# When 90% of the variance is not enough: residual EMG from muscle synergy extraction influences task performance

**DOI:** 10.1101/634758

**Authors:** Victor R. Barradas, Jason J. Kutch, Toshihiro Kawase, Yasuharu Koike, Nicolas Schweighofer

**Affiliations:** Biomedical Engineering, University of Southern California, Los Angeles, California, USA; Biokinesiology and Physical Therapy, University of Southern California, Los Angeles, California, USA; Precision and Intelligence Laboratory, Tokyo Institute of Technology, Yokohama, Japan

**Author notes:** Corresponding author (NS).

## Abstract

Muscle synergies are usually identified via dimensionality reduction techniques, such that the identified synergies reconstruct the muscle activity to a level of accuracy defined heuristically, such as 90% of the variance explained. Here, we question the assumption that the residual muscle activity not explained by the synergies is due to noise. We hypothesize instead that the residual activity is structured and can therefore influence the execution of a motor task. Young healthy subjects performed an isometric reaching task in which surface electromyography of 10 arm muscles was mapped onto estimated two-dimensional forces used to control a cursor. Three to five synergies were extracted to account for 90% of the variance explained. We then altered the muscle-force mapping via “hard” and “easy” virtual surgeries. Whereas in both surgeries the forces associated with synergies spanned the same single dimension of the virtual environment, the muscle-force mapping was as close as possible to the initial mapping in the easy surgery and as far as possible in the hard surgery. This design therefore maximized potential differences in reaching errors attributable to the residual muscle activity. Results show that the easy surgery produced much smaller directional errors than the hard task. In addition, systematic estimations of the errors for easy and hard surgeries constructed with 1 to 10 synergies show that the errors differ significantly for up to 8 synergies, which account for 98% of the variance on average. Our study therefore indicates the need for cautious interpretations of results derived from synergy extraction techniques based on heuristics with lenient levels of accuracy.

**Author summary:** The muscle synergy hypothesis states that the central nervous system simplifies motor control by grouping muscles that share common functions into modules called muscle synergies. Current techniques use unsupervised dimensionality reduction algorithms to identify these synergies. However, these techniques rely on arbitrary criteria to determine the number of synergies, which is actually unknown. An example of such criteria is that the identified synergies must be able to reconstruct the measured muscle activity to at least a 90% level of accuracy. Thus, the residual muscle activity, the remaining 10% of the muscle activity, is often disregarded as noise. We show that residual muscle activity following muscle synergy identification has a large systematic effect on movements even when the number of synergies approaches the number of muscles. This suggests that current synergy extraction techniques may discard a component of muscle activity that is important for motor control. Therefore, current synergy extraction techniques must be updated to identify true physiological synergies.

## Introduction

One of the most salient problems the central nervous system (CNS) faces when generating movements is the redundancy of the motor system [1]. That is, the CNS can generate an infinity of different motor commands to produce the same action. This redundancy spans the length of the causal chain of motor control: from neuron to muscle to joint levels. In light of the complexity of this problem, the muscle synergy hypothesis posits that the CNS groups the control of functionally similar muscles into modules called muscle synergies [2]. This would reduce the number of variables that the CNS needs to control to produce a movement, decreasing the complexity of the computations necessary for motor control [3].

Direct evidence for the muscle synergy hypothesis comes from experiments in animal models [3–6]. These show that simultaneous stimulation of different groups of motor neurons elicits movements that correspond to the superposition of the movements obtained by stimulating each group of neurons separately [3, 5, 6]. However, most of the supporting evidence in humans is indirect and comes from measurements of electromyography (EMG) from multiple muscles during a variety of motor tasks [7–11]. Dimensionality reduction techniques, such as non-negative matrix factorization, show that different muscles tend to co-activate in reliable patterns during task execution [12]. One of the interpretations of these results is that they reveal the grouping of muscles into functional synergies [8, 9, 13, 14]. An alternative interpretation, however, is that the discovered patterns arise because of biomechanical constraints imposed by the task [15–17].

This controversy notwithstanding [18], dimensionality reduction techniques for the extraction of muscle synergies rely on the ability of the extracted synergies to reconstruct the originally measured EMG signals accurately [19]. That is, the extracted synergies must capture a high proportion of the variability in the recorded EMG, attributing the discarded or residual variability in the data to measurement and process noise. This proportion is usually adjusted by making the number of muscle synergies a hyper-parameter to be tuned to best fit the data [20]. A widely used rule of thumb is to set the number of muscle synergies to the minimum number that accounts for at least 90% of the variability in the EMG.

However, this method neglects the fundamental role of muscle synergies as building blocks of movement, as the ability of the extracted muscle synergies to reconstruct the observed movement is often ignored [19, 21, 22]. Indeed, the ability of muscle synergies to reconstruct measured forces in an isometric task at the wrist becomes largely degraded as the number of considered muscle synergies decreases [22]. This is true even when the extracted synergies capture an acceptable portion of the variability in the EMG signals according to the defined heuristics. This suggests that the portion of EMG variability that is not captured by the extracted muscle synergies is important for a full description of the motor action.

Here, we therefore aimed to determine the importance of the residual EMG in the execution of a motor task. We tested the null hypothesis that following extraction of muscle synergies with non-negative matrix factorization and using the 90% of explained variance rule to select the number of synergies, the residual muscle activity is due to noise. Therefore, if our experimental data failed to support this hypothesis, it would suggest that the residuals are structured and can therefore influence motor performance.

To this end, we used the virtual surgery paradigm, which simulates tendon transfer surgeries [10]. The virtual surgery alters the pulling forces of arm muscles in a virtual mapping from EMG to two-dimensional isometric force at the wrist, which affects performance during the reaching task. This EMG-force mapping can be simplified into a synergy-force mapping by combining the pulling forces for each arm muscle according to a set of previously identified muscle synergies. Given that the number of muscles is necessarily larger than the number of extracted synergies, it is possible to build virtual surgeries that produce identical synergy-force mappings but different EMG-force mappings. We exploited this property by designing virtual surgeries that modified the EMG-force mapping to two opposite extremes while producing the same synergy-force mapping.

The “easy” surgery modified the EMG-force mapping as little as possible with respect to the baseline mapping, and the “hard” surgery modified the mapping as much as possible. The two virtual surgeries were designed based on the extracted muscle synergies that account for at least 90% of the variability in the EMG. Consequently, the effect of the surgery on the residual portion of the EMG was not specified, leading to possible differences in the effects of the easy and hard surgeries on task variables. If the EMG residuals are attributable to noise, then both surgeries should produce similar errors in the direction of reaching when introduced suddenly. Alternatively, if the EMG residuals have a latent structure, then both surgeries should have a differential effect on the residuals and on the error in the direction of reaching. We found that the sudden introduction of both kinds of virtual surgeries produced largely different errors, supporting the existence of a latent structure in the EMG residuals.

## Materials and Methods

### Subjects

Fifteen right-handed subjects (mean age, 27.9 ± 8.75 years, s.d.; thirteen males) participated in the study after providing written informed consent. All procedures were approved by the Ethical Review Board of the Tokyo Institute of Technology.

### Experimental setup

Each participant sat on a racecar seat while gripping a handle located at the height of the base of their sternum with their right hand. The arm posture corresponded to an elbow flexion of around 90° and the elbow was supported on a stand at approximately the same height as the hand. A splint was used to immobilize the hand, wrist and forearm. Participants were instructed to lean on the back of the seat for the duration of the experiment. The base of the handle was attached to a six axis force transducer (Dyn Pick; Wacoh-Tech Inc.) used to measure isometric forces. The force transducer was mounted on a 2-D sliding rail to allow for an adjustable configuration for each participant. A virtual environment was displayed on a computer screen placed at the height of the participants’ eyes at a distance of around 1 m. The virtual environment consisted of a circular red cursor (1 cm diameter), and several ring-shaped white targets (2 cm diameter) on a black background.

We recorded surface EMG activity from 10 muscles crossing the shoulder and elbow joints: pronator teres, brachioradialis, biceps brachii long head, triceps brachii lateral head, triceps brachii long head, anterior deltoid, middle deltoid, posterior deltoid, pectoralis major, and middle trapezius. Active bipolar electrodes (DE 2.1; Delsys) were used to record EMG activity. EMG signals were bandpass filtered (20-450 Hz) and amplified (gain 1000, Bagnoli-16; Delsys). Force and EMG recordings were digitized at 2 kHz using an USB analog-to-digital converter (USB-6259; National Instruments).

To reduce random oscillations of the cursor caused by the stochastic nature of EMG signals, a mass-spring-damper dynamics filtered the EMG signals further [10]. The mass-spring-damper dynamics governed the movement of the cursor according to:

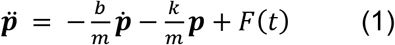

where ***p*** is a vector containing the x and y positions of the cursor on the screen and its derivatives are indicated in dot notation, m is the system’s mass, k is the stiffness, and b is the damping coefficient (m = 0.05 kg, b = 100 kg/s). F(t) is the force recorded by the force transducer (during force control) or the estimated force by the EMG-force mapping (during EMG control). k was calculated as a function of the maximum voluntary force (MVF) (described in the next section), so that targets at equal percentages of MVF required the same cursor displacement across participants.

### Experimental protocol

In all phases of the experiment, participants performed isometric force tasks. These tasks required the displacement of a cursor on a visual display from a center position to one of eight targets radially and uniformly distributed around the center. Participants first performed a force control task and then an EMG control task (Fig 1a). In the force control task, the cursor was controlled via forces applied by the arm on a load cell (force control). In the EMG control task, the cursor was controlled by a linear approximation of the force derived from EMG measurements of 10 arm muscles (EMG control).

**Fig 1.**
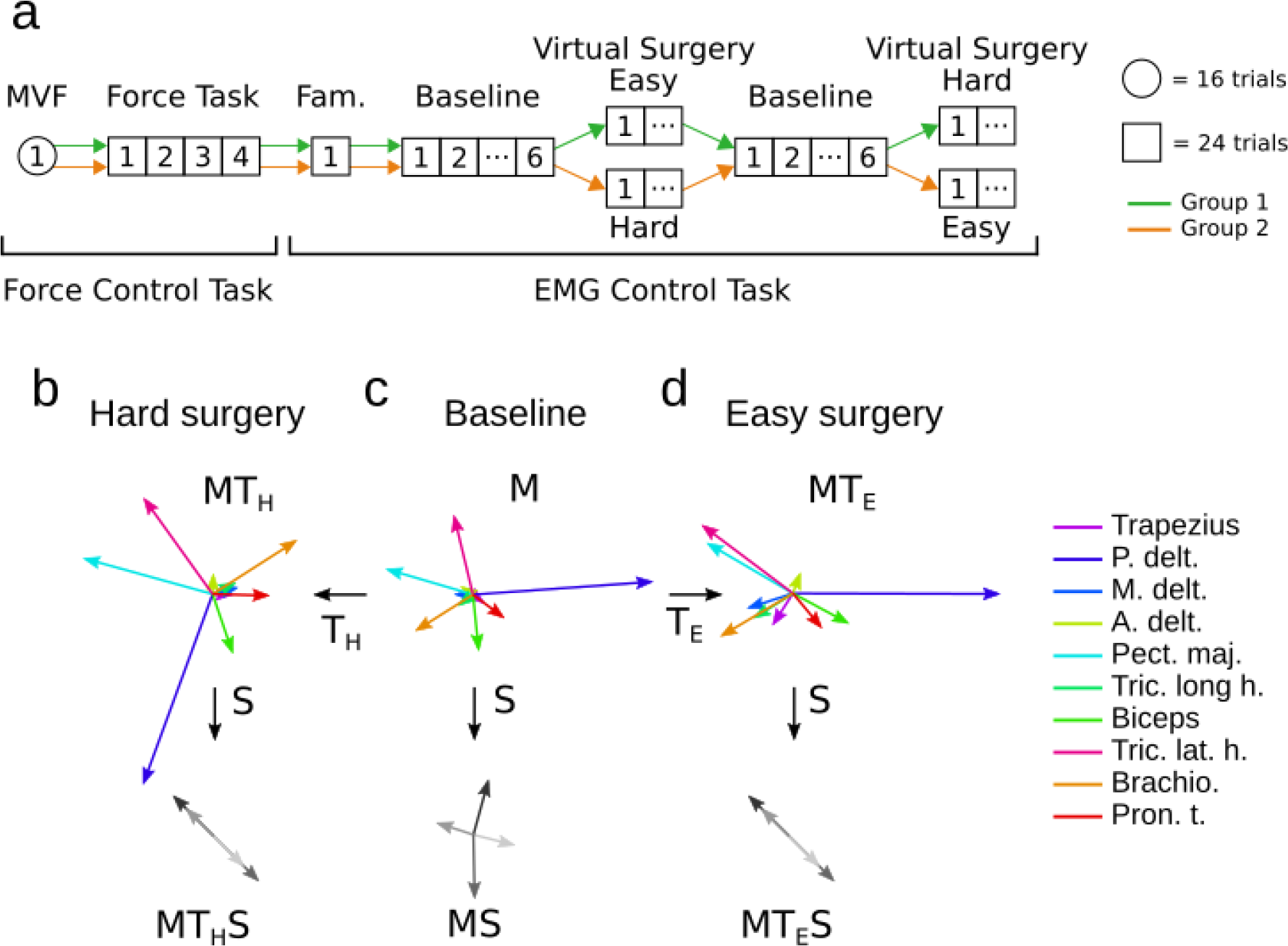
Experimental schedule and virtual surgery construction. **a**. Experimental schedule. Following a maximum voluntary force (MVF) block, participants performed a force control task. Simultaneous recording of EMG and force data in this task were analyzed to extract muscle synergies, produce the baseline EMG-force mapping, and construct the easy and hard incompatible virtual surgeries. Participants then performed the EMG control tasks, starting with a familiarization block, followed by baseline, and then one of the two virtual surgeries (easy or hard). In this cross-over study, participants then performed the other virtual surgery following a new baseline. Note that in this study, we only analyzed the data from the first block of each of the two virtual surgery procedures. **b - d**. Virtual surgery construction. **c**. EMG-force mapping extracted after the force control task for one participant. Each arrow represents the estimated force on the horizontal plane that a single muscle would produce when fully activated in isolation from the rest of the muscles (columns of M matrix). Forces produced by each of the muscle synergies extracted after the force control task (columns of MW matrix). Before applying any virtual surgery, these forces span the 2-dimensional plane completely. **b**. Hard incompatible surgery. We designed an incompatible surgery by rotating the force vectors in MS so that they became collinear at an angle of 135° degrees while maximizing the angles between the column vectors of M and MT_H_ while producing the desired MT_H_S. **d**. Easy incompatible surgery. We obtained the easy incompatible surgery by minimizing the angles between the column vectors of MT and MT_E_ while making MT_E_S equal to MT_H_S. This way the individual synergies produced the same force in both cases.

The force control task started with a maximum voluntary force (MVF) block, in which participants were instructed to produce a maximum voluntary force with their right arm in each of eight directions spanning the horizontal plane, with two trials for each direction. The mean MVF was calculated as the mean of the maximum forces recorded across all trials. For each muscle, the value at the 95 percentile of the recorded EMG signal across all trials was used to normalize the values of EMG from the corresponding muscle in all subsequent tasks.

Participants then performed an isometric reaching task by applying force with their right arm to reach targets in the virtual environment. The recorded force and EMG signals during this task were processed to compute the EMG-force mapping, extract muscle synergies, and construct the virtual surgeries. Targets were arranged radially in eight directions and required 5, 10, 15 or 20% of MVF to be reached. Each trial started by displaying the target at the central position. The central position corresponded to the position of the cursor when no forces were applied. After placing the cursor inside the central target for two seconds, the central target disappeared and one of the radial targets appeared. After reaching each target, both the cursor and the target disappeared from the screen and participants were asked to hold the applied force as steadily as possible for two seconds. Next, the cursor and the central target reappeared and participants were asked to move the cursor back to the center. After this, another trial began. Each target was presented three times, with a total of 96 trials. Targets were presented in a randomized order. Trials were repeated if participants failed to reach a target.

Next, cursor control was switched to EMG control without the knowledge of the participants, after which participants performed the reaching task under EMG control. The first EMG control block was a familiarization block, and was followed by one type of incompatible surgery, easy or hard, followed by the other in a cross-over design (see Fig 1A). The order of the easy and hard surgeries was pseudo-randomized such that 7 participants started with the easy surgery. Participants rested for 5 minutes between surgery types. Each surgery condition consisted of three phases: baseline, virtual surgery, and washout, which consisted of 6, 12, and 6 blocks, respectively. Each block consisted of 24 trials: three trials for each of the eight targets at a magnitude of 10% MVF randomized within target sets containing each one of the eight targets. The level of baseline noise in each EMG signal was measured at the start of every block while the participant was relaxed. This baseline noise was subtracted from the EMG signals measured during the corresponding block.

Note that in this study, we focus exclusively on data recorded during the first set of eight targets following the onset of each virtual surgery. Analysis of the following blocks for each surgery will be covered in a separate manuscript that focuses on learning of incompatible virtual surgeries.

### EMG-force mapping

Force produced at the hand with the arm in a static posture can be approximated as a linear function of the activations of muscles that actuate the shoulder and elbow [10]:

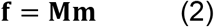

where **f** is a two-dimensional force vector produced on the horizontal plane, **m** is a ten-dimensional vector of muscle activations, composed by normalized EMG signals recorded from ten muscles simultaneously, and **M** is a 2 × 10 matrix that maps muscle activations to forces. **M** was determined via linear regression of 10 EMG signals against 2D forces recorded during every trial of the main force control subtask. Before performing the regression, forces were low-pass filtered (second-order Butterworth; 1 Hz cutoff) and EMG signals were band-pass filtered (second-order Butterworth; 5-20 Hz), rectified, and normalized. The signals were recorded from the time of target go to the end of target hold.

### Synergy extraction and number of synergies

We used non-negative matrix factorization (NMF) to extract muscle synergies from the EMG signals collected during the main force control subtask:

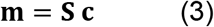

where **S** is a 10 × *N* matrix that contains the identified synergies in its columns with *N* being the number of synergies, and **c** is an *N*-dimensional vector of synergy activations. Equation 3 assumes perfect matrix factorization (no residual EMG activity).

EMG signals collected during the main force control subtask were processed in the same way as described in the EMG-force mapping section. The synergy extraction procedure closely followed a method previously described [10]. Synergies were extracted for all *N* from 1 to 10. For each case, the synergy extraction algorithm was run 100 times, and the result with the highest reconstruction quality *R*^*2*^ of the original EMG signals was kept. Two criteria were required to select *N*. The first was to set *N* as the minimum number of synergies necessary to explain at least 90% of the EMG data variance. The second involved calculating the changes in slope in the *R*^*2*^ curve as a function of *N*. Linear regressions were performed on sections of the curve between *N* and 10. *N* was selected as the smallest value for which the mean squared error of the linear regression was < 10^−4^ [11]. If the two criteria did not match, *N* was selected as the case in which the extracted synergies had the smallest number of similar preferred directions (number of adjacent directions separated by less than 20°). This occurred for seven of the participants.

### Construction of easy and hard incompatible surgeries

As in a previous study [10], we designed virtual surgeries that were incompatible with the muscle synergies extracted by nonnegative matrix factorization (NMF) [23]. A virtual surgery modifies the EMG-force mapping (**M**) by applying a linear transformation in muscle space [10]:

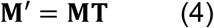

where **T** is a 10 × 10 matrix that constitutes the transformation or virtual surgery.

Incompatible virtual surgeries are designed such that muscle activations **m** produced by synergy combinations **Sc** are restricted to generate forces along only one dimension of the force space, while the resulting EMG-force mapping **M’** spans the whole force space. Therefore, theoretically, any force can still be produced by a new combination of muscle activations **m’**, but in practice, produced forces are biased towards one dimension of the plane.

It is important to note that the set of incompatible surgeries is infinite. This is because the number of muscles used in the virtual mapping is larger than the number of muscle activity patterns found using muscle synergy analysis. A previous study [10] combined randomness and difficulty matching to select compatible and incompatible virtual surgeries.

In contrast, here we specified a series of constraints to yield only two possible virtual surgeries. Specifically, we built hard **T**_**H**_ and easy **T**_**E**_ incompatible surgeries such that they were equivalent in the force space spanned by each participant’s extracted muscle synergies (Figs 1b and 1d). We first note that according to equations 2, 3 and 4, forces produced during the surgery are given by:

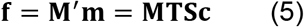

assuming that muscle activations are generated by combinations of synergies. This equation shows that surgery **T** can alternatively be thought to transform the extracted synergies **S** into a new set of synergies:

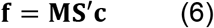

In order to build an incompatible surgery it is necessary to find **S’** such that the matrix **MS’** is rank deficient. This guarantees that forces produced by this mapping lie in a single dimension. Geometrically, this means that the forces associated with each individual synergy from **S’** are collinear (Figs 1c and 1g).

Easy surgeries were built such that the angles between the column vectors of the original **M** mapping and of the transformed mapping **M’** were as small as possible. In contrast, hard surgeries were built by making these angles as large as possible (Fig 1b). These conditions produced **M’** mappings that are similar or very different to the original **M** mapping in the case of easy or hard surgeries, respectively. For this, we used a two-step optimization procedure to first obtain a transformed set of synergies **S’**, and second, to compute the incompatible surgery **T**. We constrained **S’** to be equal for both the easy and hard incompatible surgeries. This ensured that the only difference between both virtual surgeries is the transformed mapping **MT**. We chose a configuration such that the individual force vectors associated to each synergy in **S** were rotated onto a line that bisected the plane at an angle of 135° with the x-axis. Therefore, each force vector conserved its magnitude, and its direction was assigned to the direction of the bisecting line that was closest to it: 135° or −45°. This can be represented as a system of equations in which the elements of **S’** are the unknowns:

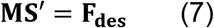

where **F**_**des**_ is a 2 × *N* matrix containing the desired force components associated with each synergy after the virtual surgery in each of its columns, with *N* being the number of extracted synergies. This problem has 10*N* unknowns and only 2*N* equations, so we introduced an optimization objective to arrive to a unique solution. A reasonable objective is to minimize the sum of the squares of the elements of **S’**, as this creates a sparse set of synergies. Additionally the elements of **S** are required to be non-negative. This optimization problem can be posed as a quadratic program:

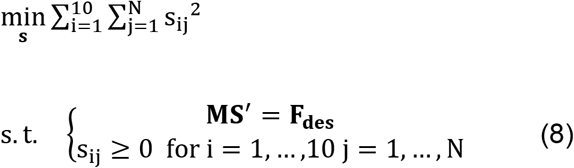

We transcribed this quadratic program into its canonical form and solved it using the *quadprog* function in Matlab.

After obtaining **S’**, we computed the incompatible surgery **T** by noting that

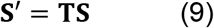

This is a system of equations where the elements of **T** are the unknowns. We note that **T** is a 10 × 10 matrix, so in this case there are 100 unknowns and 10 *N* equations. The system is overdetermined in all cases where *N* < 10, which in our case is guaranteed.

In order to find the easy virtual surgery, we used our requirements of similarity between **M** and **M’** to introduce an optimization objective to arrive to a unique solution. **M** and **M’** are considered similar when the angles between their corresponding column vectors are as small as possible. The cosine of the angle between two vectors is proportional to the dot product of both vectors.

Therefore, we defined the optimization objective as

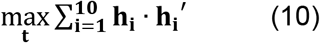

where **h**_**i**_ and **h**_**i**_’ are the column vectors of **M** and **M’**, respectively. This optimization objective is not bounded, so we added constraints to the magnitude of the resulting **h’** vectors:

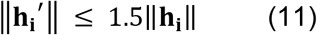

This problem can be posed as a linear program with quadratic constraints, with equation 10 as the objective, and equations 9 and 11 as equality and inequality constraints, respectively. The result of this optimization procedure yields **T**_**E**_, the easy incompatible virtual surgery.

In order to compute the hard incompatible virtual surgery **T**_**H**_, the procedure is the same as for the easy incompatible surgery. The only difference is that the optimization objective is minimized instead of being maximized. In turn, this maximizes the angles between **h**_**i**_ and **h**_**i**_’:

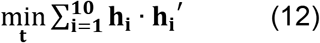

Both linear programs with quadratic constraints were solved using the *fmincon* function in Matlab.

## Data analysis

### Task performance metric

We used the initial angular error as a metric to quantify task performance during the experiment, before possible feedback corrections. The initial angular error was calculated for each trial as |θ_target_ − θ_cursor_|. θ_target_ is the direction of the target. θ_cursor_ is defined as the direction of the line segment that joins the point at which the cursor exits a 2 cm diameter circumference at the center of the screen and the position of the cursor 100 ms after exiting the circumference. We averaged the initial angular error for the targets within sets of eight trials. We only took into account targets that were not aligned with the line of action of the surgery. That is, targets other than those at 135° and −45° from the horizontal on the screen.

### EMG residual analysis

We analyzed the residual EMG signals obtained after reconstructing the measured EMG signals based on the extracted muscle synergies. After synergy extraction using the NMF algorithm, and extending equation 3, muscle activations can be represented as

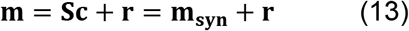

where **m**_**syn**_ is the synergy component of muscle activation, and **r** is the residual component of muscle activation that cannot be accounted for by the extracted synergies. Consequently, the forces associated with the EMG signals by the EMG-force mapping have a synergy and a residual component:

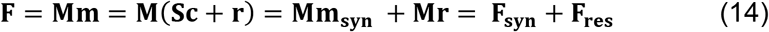

where **F**_**syn**_ and **F**_**res**_ are the synergy and residual components of force, respectively. Because virtual surgeries are built based on **S**, the intended effects of the virtual surgeries are only manifested on the synergy component of force, and the effect on the residual force is not specified.

In order to decompose a given EMG sample **m** into its synergy and residual components (**m**_**syn**_ and **r**), we first computed **m**_**syn**_ via non-negative least squared regression of **S** and **m**, which yielded **c**. This algorithm optimizes the same cost function as the NMF algorithm. Therefore, using equation 13, **m**_**syn**_ is given by the product of **S** and **c**. Consequently, **r** is found by subtracting **m**_**syn**_ from **m**.

We then analyzed the effects of the surgery on both the synergy and residual components of EMG. For this, we used the EMG activity that participants produced when they acquired each target during the first baseline phase of the experiment. We then separated the average EMG activity of each subject and target **m** into **m**_**syn**_ and **r**.

We also estimated both force components **F**_**syn**_ and **F**_**res**_ produced for each target at the onset of the easy and hard virtual surgeries by substituting **M** by **M’** in equation 14. We then compared the estimated force direction to the intended direction for each target to obtain an estimate of the error that subjects would produce at the onset of each virtual surgery.

### Shuffling of EMG residuals

Shuffling the residual component of different EMG signal samples creates random residual components with the same statistical properties as the original residuals. If the residual EMG activity can be disregarded as noise, then shuffling the residuals should have no significant effect on the estimated forces with respect to pre-shuffling. On the contrary, if the residuals have a structured organization, shuffling the residuals would destroy this organization. Consequently, the force estimates would most likely be different from the pre-shuffling estimates. We therefore shuffled the residual components of the EMG samples that we used to estimate forces, and re-estimated the total forces that would be produced at the onset of the easy and hard virtual surgeries. We averaged the results of 1000 different shuffling instances.

## Results

### Hard incompatible virtual surgeries produced larger initial angular errors than easy incompatible virtual surgeries

The number of extracted muscle synergies *N* for all subjects ranged from three to five (*N* = 3, 1 subjects; *N* = 4, 11 subjects; *N* = 5, 3 subjects). Fig 2 shows sample cursor trajectories before and after the onset of the virtual surgeries. Both surgeries produced a bias in the cursor movement along the designed direction as predicted, although cursor movements were not perfectly constrained to this direction. Overall, deviations from the line of action of the surgery were closer to the intended target during the easy surgery than during the hard surgery (Fig 2b).

**Fig 2.**
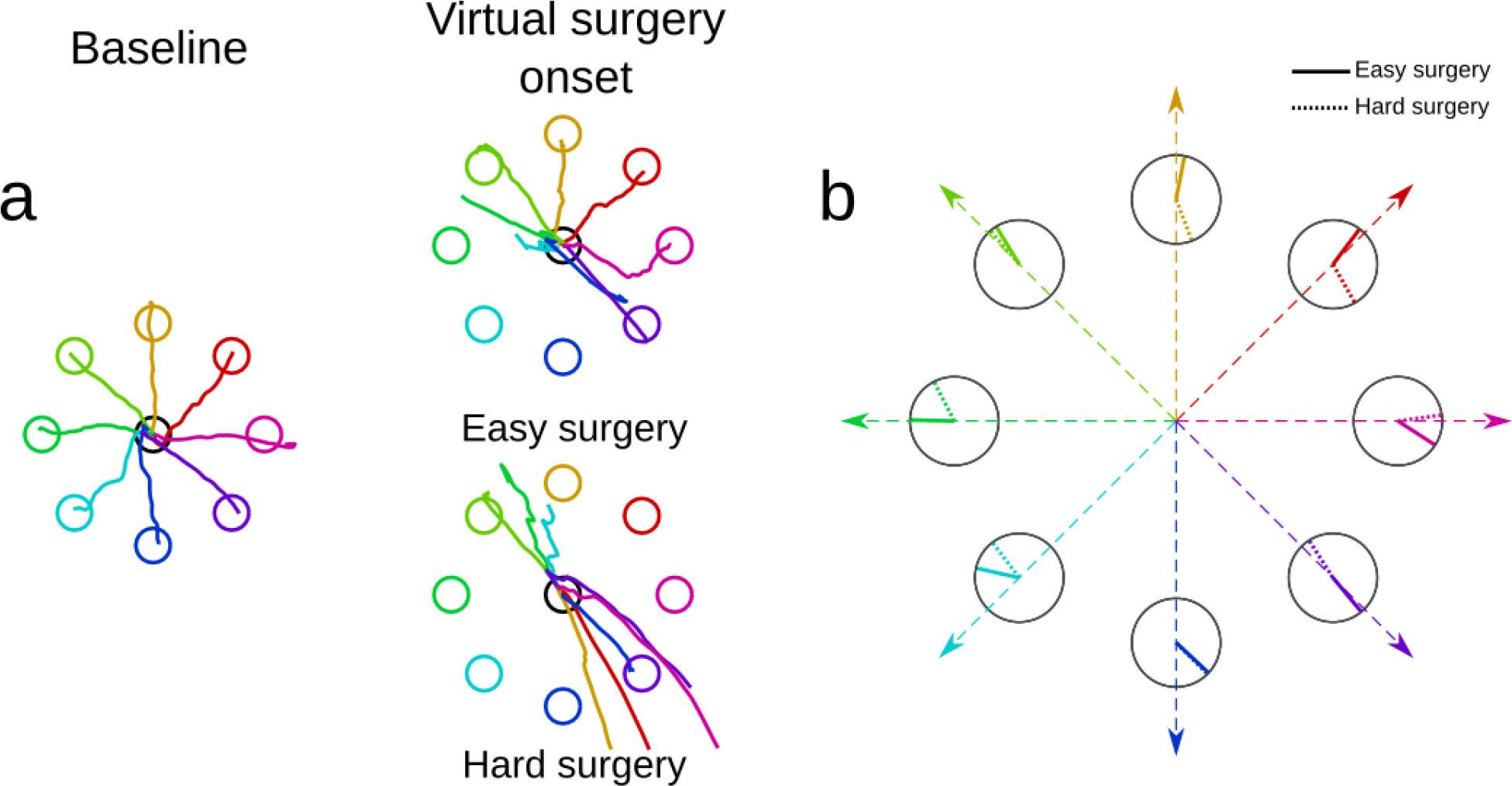
Example of cursor trajectories during the EMG control task by a representative subject. **a**. Sample cursor trajectories. These trajectories correspond to the last target set of the baseline subtask, and the first target set after the onset of the hard and easy incompatible virtual surgery tasks. This subject experienced the easy virtual surgery first. The trajectories tended to fall along the line of action of the virtual surgery, notably in the hard surgery. **b**. Comparison of initial directions of cursor movement between the onset of the easy and hard virtual surgeries. Straight-line segments represent the computed direction of movement of the cursor depicted in panel a 100 ms after exiting the central position. Solid lines correspond to the initial directions during the easy surgery onset and dotted lines correspond to the hard surgery onset. This subject produced larger initial errors at the onset of the hard virtual surgery than at the onset of the easy surgery (see targets at 45° and 90°).

Over all 15 participants, the mean error for the first set of targets after the onset of the surgery was clearly larger for the hard surgery than for the easy surgery (hard surgery: 81.4° ± 3.8° s.e, easy surgery: 54.5° ± 4.6° s.e., p < 10^−3^, paired t-test; see Fig 3B, experiment). This difference in errors may appear surprising at first, given that the easy and hard surgeries had the same effect on the synergy component of the force. That is, they restricted the forces associated with the synergies along one dimension. However, the synergies were only required to account for 90% of the variance in EMG. Therefore, the EMG residuals appeared to generate an additional component of force.

**Fig 3.**
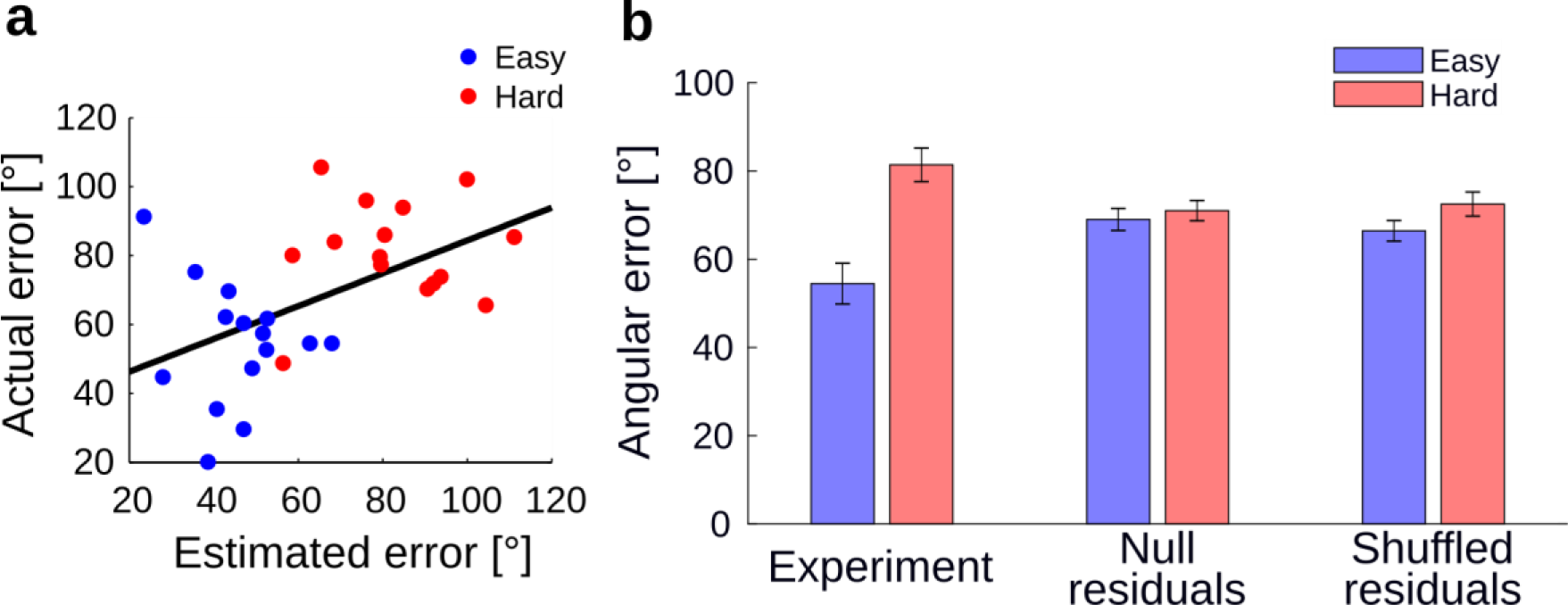
Comparison between actual directional cursor errors and estimated errors. **a**. Correlation between the estimated average angular error after applying the surgery and the actual average error during the first target set of the virtual surgery for all participants. **b.** Actual and estimated errors in initial direction of force at the onset of the easy and hard virtual surgeries. In the experiment, the hard surgery produced a larger initial error than the easy surgery. As a comparison, errors were also estimated: i) using only the synergy component of the average EMG signals (null residuals), and ii) using the shuffled residual component of the EMG. The null residual estimate produced error estimates that showed no difference between the hard and the easy virtual surgeries. The EMG signals with shuffled residuals produced error estimates that were similar to those obtained using only the synergy component of the EMG signal.

### Initial angular error was determined by effect of surgery on residual EMG activity

To verify the effect of residuals on movement error, we decomposed the recorded EMG signals into their synergy and residual components (equation 14). Fig 4 shows the estimated forces corresponding to the total EMG activity **F** (top), and the synergy (**F**_syn_) and residual (**F**_res_) components of force (middle and bottom, respectively) at each target for a representative subject (equation 14). The center column shows **F**, **F**_**syn**_ and **F**_**res**_ before the onset of the surgeries. The left and right columns show **F**, **F**_**syn**_ and **F**_**res**_ after applying the hard and easy surgeries, respectively. The incompatible design of the surgery can be appreciated on **F**_**syn**_, as these forces lie on the 135° line of action of the virtual surgery (Fig 4, middle row).

**Fig 4.**
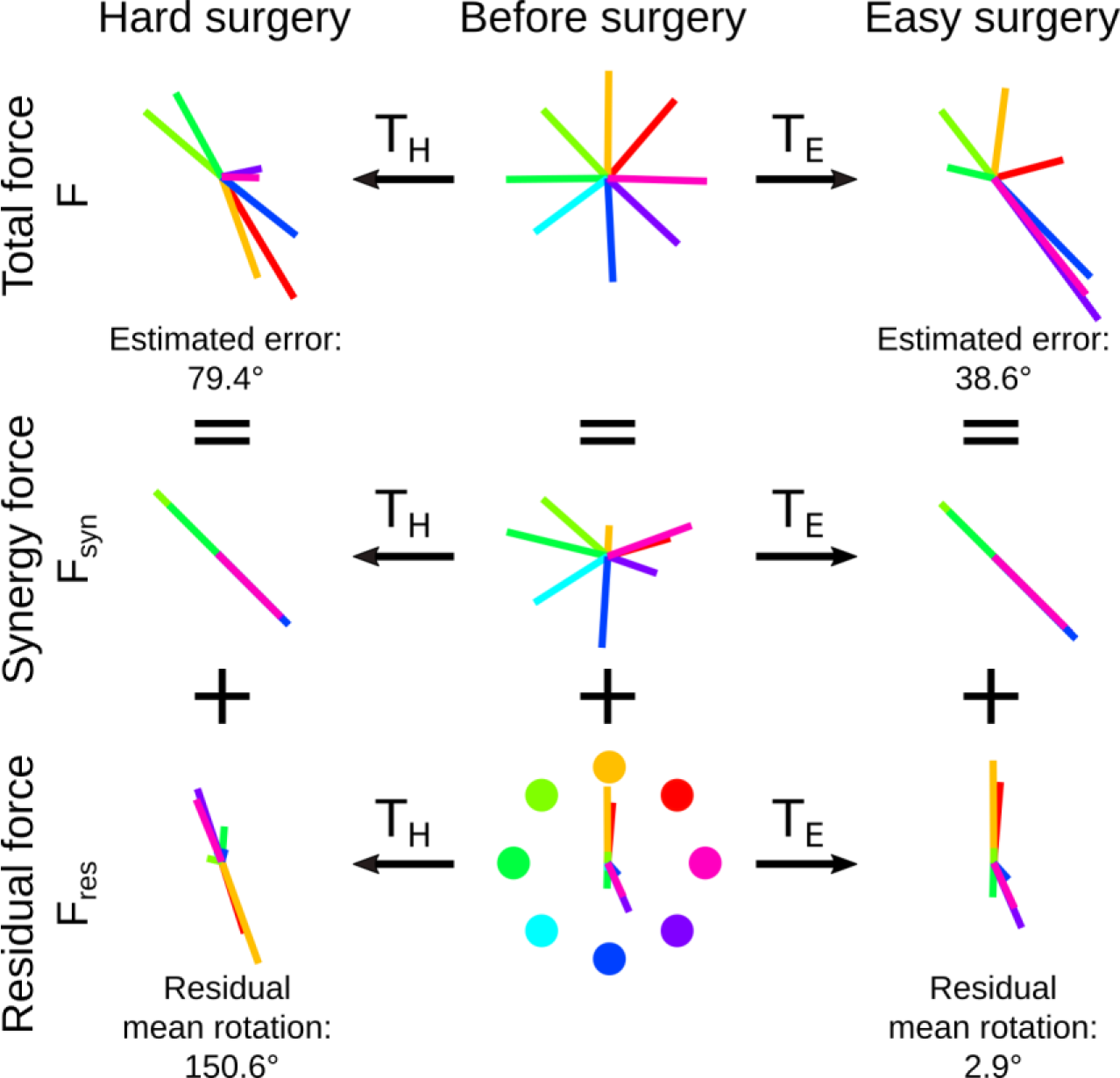
Residual forces can explain the differences in initial direction error at surgery onset between the easy and hard surgery conditions. Top row: Estimated forces **F** at each target before and after applying the hard and easy virtual surgeries. We indicate the average estimated error across targets for each virtual surgery. Middle row: Estimated synergy components of force **F**_**syn**_ at each target. Bottom row: Estimated residual components of force **F**_**res**_ at each target. We indicate the average rotation of **F**_**res**_ after each virtual surgery with respect to **F**_**res**_ before the surgery. Middle column: **F**, **F**_**syn**_ and **F**_**res**_ before the surgery. Left and right columns: **F**, **F**_**syn**_ and **F**_**res**_ after applying the hard and easy surgeries, respectively. Colors represent the eight targets in the task as indicated in the middle bottom diagram. Data shown in this figure corresponds to the same representative subject as in Fig 2.

Given that the EMG signals that we used to estimate forces were representative of the subjects’ actions during baseline, and assuming that subjects produced these EMG signals when suddenly exposed to the virtual surgeries, the directions of the estimated forces after applying the virtual surgery (Equation 5) also provided an estimate of the cursor error to each target (Fig 4, top row). These initial error estimates were consistently higher for the hard surgery than for the easy surgery (hard surgery: 82.75° ± 4.19° s.e., easy surgery: 45.57° ± 3.03° s.e., p < 10^−3^, paired t-test), and qualitatively matched the experimental results of the cursor error (robust regression, slope = 0.47 ± 0.15 s.e., p = 0.004, R^2^ = 0.25) (Fig 3a).

Errors following the easy and hard surgeries can be explained by the residual’s structure (Fig 4, bottom row). The hard surgery produced a mean rotation of **F**_**res**_ with respect to baseline that was much larger than that produced by the easy surgery (hard surgery: 113.60° ± 10.15° s.e., easy surgery: 4.42° ± 1.5° s.e., p < 10^−3^, paired t-test). Note that although we did not specify the effect of the virtual surgery on the residual component of force, we found that it is stereotypical according to the type of surgery.

### Shuffling residual EMG activity revealed structure in the residuals

We then shuffled the residual EMG components among trials to all targets to demonstrate possible structure. Initial error estimates based on the shuffled signals did not indicate a significant difference in average initial error between the easy and hard virtual surgeries (easy surgery: 66.42° ± 2.31° s.e., hard surgery: 72.41° ± 2.72° s.e., paired t-test, p = 0.10) (Fig 3b, shuffled residuals). Furthermore, the magnitude of this error lied at an intermediate level between the errors observed experimentally for the easy and hard surgeries. Importantly, the means of the estimates produced by shuffled signals were indistinguishable from estimates produced based on a null residual condition, that is, exclusively using the synergy component of the EMG to produce estimates (easy surgery, paired t-test, p = 0.45; hard surgery, paired t-test, p = 0.68) (Fig 3b, null and shuffled residuals).

### Estimated differences between errors for easy and hard surgeries remained significant for high-dimensional synergy sets

We tested whether building virtual surgeries based on synergy sets with a larger *N* would abolish the differences in initial direction error observed in the experiment. For each participant we built easy and hard surgeries based on surgeries considering *N* = 1, …, 10 and applied the newly constructed surgeries to the same set of EMG signals that we used to estimate errors after the introduction of the surgery. This allowed us to estimate the errors that participants would have produced if they had experienced these surgeries.

We found that the surgeries produced estimated differences in initial direction errors that were maximal for *N* = 1, and gradually decreased until disappearing at *N* = 10 (as expected, since activity from 10 muscles was recorded; Fig 5). The estimated error differences remained significant up to *N* = 8 (p = 0.001, paired t-test). This indicates that the residual components of EMG produced a differential effect on the estimated error even when high-dimensional synergy sets that explained a portion of the variance that largely exceeded the heuristic rule requirements (R^2^ = 0.98 ± 0.0087 s.e. at N = 8) were used to build the virtual surgeries.

**Fig 5.**
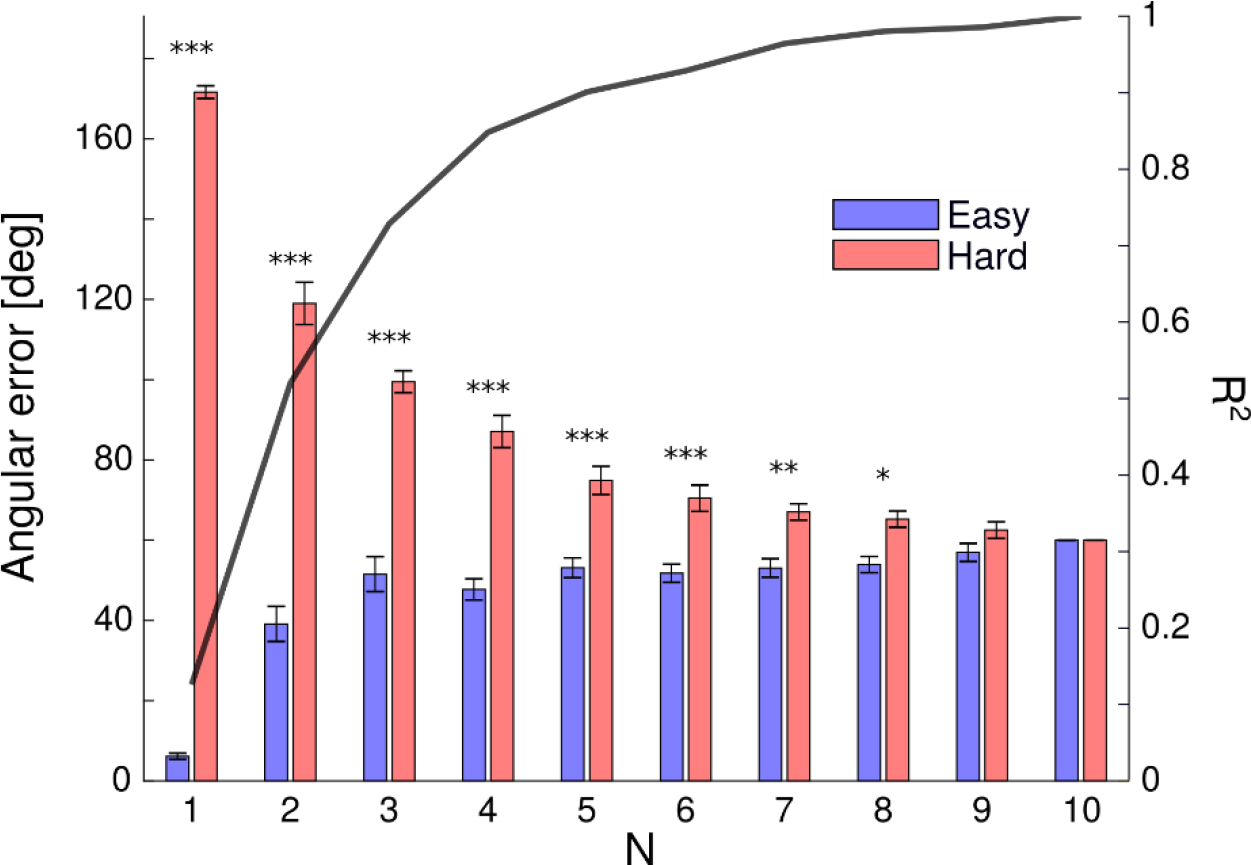
Estimated errors in initial direction when using easy and hard virtual surgeries based on muscle synergy sets with *N* = 1,…,10. Differences in estimated errors between easy and hard surgeries were significant up to *N* = 8 (p = 0.001, paired t-test). Bars represent the mean estimated error across the 15 participants and error bars represent the standard error of the estimated error. The significance of the difference between the estimated errors in the hard and easy surgeries is indicated with asterisks on top of each pair of bars. ***: p < 0.0001, **: p < 0.001, and *: p < 0.005 The solid black line shows the mean across participants of R^2^, the reconstruction quality of the baseline EMG signals used for the error estimation when considering *N* = 1,…,10.

## Discussion

Muscle synergy extraction techniques require that combinations of the identified synergies reconstruct the measured muscle activity to a heuristically defined level of accuracy, such as accounting for at least 90% of the variance in the EMG. These techniques therefore attribute the residual muscle activity not reconstructed by the identified synergies to noise. Here we studied the importance of residual EMG activity in the execution of a virtual motor task. We designed the virtual task based on a virtual surgery [10, 24] and exploited the property that virtual surgeries can produce equivalent muscle synergy-force mappings while resulting in different individual muscle-force mappings. We tested two different virtual surgeries that shared a common muscle synergy-force mapping, but differed maximally in their individual muscle-force mappings (easy and hard virtual surgeries). The surgeries had the desired effect only on the portion of the EMG signals explained by the extracted muscle synergies, defined to account for at least 90% of the variability in the signal. Therefore, the effect on the residual EMG variability was unspecified, allowing for a possible differential effect on the performance of the task.

We found that participants produced larger errors at the onset of the hard surgery than at the onset of the easy surgery. We were able to predict this result qualitatively (Fig 3a) by estimating the forces and errors that would be produced during each virtual surgery by using representative EMG signals recorded during the baseline phase of the experiment and transforming the estimated forces using the virtual surgeries. Importantly, this procedure also allowed us to separate the recorded EMG signals and the estimated forces into their synergy and residual components (Fig 4). The virtual surgeries produced the expected effects on the synergy component of the EMG. However, the easy surgery barely produced any changes on the direction of the forces associated with the residual component, whereas the hard surgery produced large changes in the direction of these forces. Given that the total force is equal to the sum of the synergy and residual components, any difference between both virtual surgeries in the estimated force and error must arise from the difference in the residual components. This provides evidence that the residual component of the EMG is essential for accounting for our experimental results, suggesting a latent structure in the residuals.

We also considered the alternative case in which the residual EMG activity is composed of noise. In this situation, we posited that there would be no differential effect of the easy and hard surgeries on the initial error, or that this effect would be small. To test this, we used the previously decomposed EMG signals and shuffled the residual components among all these EMG samples. This effectively destroyed any potential structure in the residual component, as they became randomized. We found that the easy and hard surgeries did not produce significant differential effects in the estimated initial error across participants when applied to the shuffled EMG signals. This analysis suggests that the residual component of the EMG cannot be disregarded as purely noise, and therefore demonstration of a latent structure in the residuals.

Dimensionality reduction techniques such as NMF are useful for extracting patterns from high-dimensional data sets, such as EMG recordings from multiple muscles. These techniques are usually able to extract as many patterns or synergies as individual muscles. However, there is no objective means for selecting the number of synergies of interest a priori given the exploratory nature of the analysis and the lack of a ground truth. Therefore, heuristic rules, such as selecting the number of synergies based on predefined goodness of reconstruction criteria are a common practice (i.e., reconstructing the data to a given level of accuracy, or finding an elbow in the goodness of reconstruction curve). These heuristic rules are necessarily ad hoc, and are tailored to produce useful results in the domain of the studied problem [25].

These heuristic rules ignore the role of muscle synergies in the generation of movement. That is, muscle synergy extraction has mainly focused on describing muscle activity in the input space, but has neglected the reconstruction of forces and movements in the task space [19, 21]. A number of studies have attempted the extension from input to task space in the scope of the study of synergies by incorporating task-relevant constraints, such as force reconstruction, in the dimensionality reduction procedure [13, 26]. However, in these studies, assumptions of linearity were made in the relationship between input and task spaces, that is, muscle activations and forces. Alternatively, other studies took a simulation approach by using muscle synergy activity derived experimentally as input to a computational biomechanical model to assess the goodness of the resulting movement reconstruction [27]. However, tuning of muscle activations during the simulations was necessary to obtain favorable results. Further difficulties in the use of computational biomechanical models to test for reconstruction of task space variables could stem from the difficulty of measuring EMG from all muscles involved in a movement and of building sufficiently accurate musculoskeletal models.

An alternative approach for studying the influence of extracted synergies on task-space variables consists in using a virtual isometric task, such as in this study and others [10, 22, 28]. Virtual tasks overcome the difficulty of obtaining complex biomechanical models, as the task can be defined by the experimenter. This way, the physics of the system are linear and known, and can be used in simulations in a straightforward way. A study using this approach showed that the reconstruction of isometric forces in an EMG-controlled task using muscle synergy decomposition is acceptable only when the number of synergies is equal to the number of considered muscles [22]. Otherwise, the reconstruction quality quickly degrades even when the number of synergies is derived from widely used heuristic rules [22]. Decreasing the number of synergies is associated with larger residual components of the EMG, which we showed to play an important role in task performance. Thus, our results further expand on this view, with the additional contribution of not being limited by the passive reconstruction of forces, but by directly manipulating the contribution of the residual component of the EMG to isometric force to highlight its importance in the execution of the task. These results emphasize the need of a shift within the community in the criteria used to evaluate the goodness of muscle synergies extracted through dimensionality reduction methods such as NMF.

Our results suggest that humans produce muscle activations that cannot be fully accounted for by linear combinations of low-dimensional sets of muscle synergies, as extracted via NMF. This is not in conflict with the notion that the CNS uses muscle synergies as building blocks of movement embedded in neural circuits, as shown by numerous animal studies [3–6]. Additionally, synergy control in a myoelectric task (isolating **m**_**syn**_ from **r**) has been shown to produce cursor trajectories similar to control through individual muscles, showing that synergy control may be useful for myoelectric interface applications [28]. However, this does not necessarily imply that the CNS is limited to a small number of muscles synergies structurally defined in neural circuits. It is entirely possible that the CNS readily learns and exploits a large number of task-dependent muscle synergies, which may vary from individual to individual [29]. In this sense, the role of muscle synergies could be viewed as a source of flexibility in the repertoire of possible motor commands and stability in movement execution, as opposed to a way to simplify the control of movement by limiting the number of control inputs [30].

The main limitation of our study is that our experimental design was originally conceived to test motor learning of virtual surgeries (to be presented in a follow-up manuscript). Therefore, the virtual surgery trials were presented in blocks after transitions from baseline trials. This could have induced a small amount of learning in the trials immediately following the perturbation or engaged exploratory behaviors due to the saliency of the perturbation. These undesired factors could explain why our initial error estimates do not match the experimental data more closely. A more appropriate design would randomly introduce catch trials for each surgery type among baseline trials to reduce learning effects. Nonetheless, because the observed differential effect between both virtual surgeries was large and robust, the results of a study that addresses these limitations would probably not be very different from our current results.

Overall, our results indicate that current muscle synergy identification techniques wrongly attribute the fraction of unexplained variability in the EMG signals to noise. Our study is not able to discern whether the structure of the residual component of the EMG is due to the inadequacy of an additive linear model of muscle synergies, additional muscle synergies left out of the analysis by the 90% variance or other heuristic rules, or due to other possible sources like individual muscle control. However, it is clear that studies that aim to infer neural structures through EMG recordings should carefully consider the role of the residual component of the EMG signals.

